# Genomic profiling of cancerous patients identifies FPR2 as an alternative immunotherapeutic target in glioblastoma multiforme

**DOI:** 10.1101/2020.12.26.424414

**Authors:** Yuqing Yang, Ting Sun, Chuchen Qiu, Dongjing Chen, You Wu

## Abstract

**Background:** Glioblastoma multiforme (GBM) is a type of high-grade brain tumor known for its proliferative, invasive property, and low survival rate. Recently, with the advancement in therapeutics for tumors such as targeted therapy, individual cancer-specific biomarkers could be recognized as targets for curative purposes. This study identified six differentially expressed genes that have shown significant implications in clinical field, including FPR2, VEGFA, SERPINA1, SOX2, PBK, and ITGB3. FPR2 was of the same protein family with FPR1, and the latter has been repeatedly reported to promote motility and invasiveness of multiple tumor forms.

**Methods:** The gene expression profiling of 40 GBM samples and five normal samples from the TCGA database were comprehensively analyzed. The differentially expressed genes (DEGs) were identified using R package and screened by enrichment analysis and examination of protein–protein interaction networks, in order to further explore the functions of DEGs with the highest association with clinical traits and to find hub genes. A qRT-PCR and Western blots were conducted to verify the results of this study.

**Results:** Our investigation showed that FPR2, VEGFA, SERPINA1, SOX2, PBK, and ITGB3 were significantly up-regulated in GBM primary tumor compared to the control group. Functional enrichment analysis of the DEGs demonstrated that biological functions related to immune systems, cell division and cell cycle were significantly increased, which were closely related to tumor progression and development. Downstream construction of PPI network analysis indicated that FPR2 was a hub gene involved in high level of interaction with CR3 and VEGFA, which played a key role in inflammatory pathways and cellular dysfunction.

**Conclusion:** FPR2, VEGFA, SERPINA1, SOX2, PBK, and ITGB3 were significantly over-expressed in primary tumor samples of GBM patients and were involved in cellular functions and pathways contributing to tumor progression. Out of these six pivotal genes, we intensively focused on FPR2, and our analysis and experimental data both suggested its efficacy as a potential biomarker, serving as an alternative immunotherapeutic target for glioblastoma multiforme.

## INTRODUCTION

Glioblastoma multiforme (GBM), one of the most ubiquitous brain tumors in adults [50], which arises from genetic alterations, is known for its lethality, aggressiveness, and high recurrence properties [24]. Based on the histopathological and clinical categorization, GBM is deemed as the Group IV glioma [9] with the most malignancy and mortality, accounting for 60-70% of all gliomas [30] and afflicting 3.19 people per 100,000 in the United States [30]. Multiple treatment methods have developed within the past few decades, including traditional methods such as surgery and radiotherapy [44] and more recent methods such as tumor electric field therapy (TTF). Nonetheless, while GBM treatments have advanced, limitations and availability of these options often associate with poor prognosis and yield unsatisfactory results [13]. The current average survival rate of GBM patients is within 15 months of diagnosis [3], and less than 5% have a 5-year survival rate [40]. Difficulty in achieving full removal of tumors as tumor cells has been encountered since GBM disseminates within the brain, which also significantly diminishes the efficacy of surgery and radiotherapy [37, 41], even though sub-total resection has associated with increasing survival [5]. TTF combination therapy also incurs a prohibitive monthly treatment cost of $20,000 for many GBM patients [4].

Genomes-driven systematic approach assisted by the increasing development of bioinformatics tools and availability of online databases, has facilitated the discovery of many molecular biomarkers in recent years. These cancer-related databases, such as The Cancer Genome Atlas (TCGA) [7–8, 35, 39], are in large-scale. Sizable numbers of oncogene microarray results can be traced from these online databases and processed statistically to bring new insights into cellular immunotherapy and molecular targeted therapy [32, 44]. Investigations into cancer-specific genomic features can reveal functional changes, transcriptomic and proteomic alternations, and molecular subtypes underlying the transformation from normal tissue to cancerous tumors.

In this study, we utilized computational tools and bioinformatics analysis, focusing intensively on comparing normal group and GBM patients, who experienced the most aggressive form of gliomas, placing statistical analysis in context. Six significant genes from the differentially expressed genes (DEGs) are selected and regarded as potential key genes to GBM formation. Our analysis eventually yielded FPR2 as the hub gene, the main focus of this study, and its importance is validated via discussion of published literature and western blotting. In summary, we provide insights for GBM treatment strategies by recognizing a novel and potential prognostic biomarker as an immunotherapeutic target, setting a realizable and promising trajectory for the future development of GBM treatment.

## MATERIALS AND METHODS

### Cell recovery and culture

GBM cells (U-87 MG) and HEB cells (normal brain glia cells) were purchased from SHANGHAI GUANDAO BIOSYNTHESIS BIOTECHNOLOGY CO., LTD. Removing and discarding culture medium, cell layer was rinsed with 0.25% (w/v) Trypsin-0.53mM EDTA solution and chelated using 3.0 ml of Trypsin-EDTA solution. Then, cell lines were treated with 0.25% trypsin/EDTA for 2-3 minutes at 37 °C and subcultured in a ratio of 1:10. All cell types were kept at 37 °C in 5% CO_2_ atmosphere and 95% humidity.

### Mining Microarray Data

The Cancer Genome Atlas (TCGA) database is an integrative collection of genomic, epigenomic, transcriptomic, and proteomic information spanning across 33 types of cancer composing of over 20,000 samples. mRNA sequencing data of glioblastoma multiform (TCGA-GBM) primary tumor sample and normal blood-derived samples were collected from the TCGA database (https://portal.gdc.cancer.gov/). A total of 40 WHO IV samples and 5 normal blood-derived samples were involved in the analysis. The age of all samples is between 40 years and 70 years of age at the time of diagnosis.

### Screening and selection of differentially expressed genes

Raw data is utilized for differential expression analysis between transcriptome profiling data of blood-derived normal samples and primary tumor tissue patients. A correlation plot was first conducted to perform the correlation among all samples to exclude the potential outliers. DEGs between samples was detected by using the DESeq2 package in R [15]. The DEGs with p-adjusted value (adjusted using BH method) < 0.05 and log2 Fold Change >1 were identified as up-regulated, while those with log2 Fold Change <−1 were identified as down-regulated. Then the result from DEG analysis was visualized using a volcano plot, heatmap hierarchical clustering, and Pearson’s correlation plot. Moreover, a box plot comparing the differential expression level of a particular gene was implemented.

### Functional enrichment and sub-networks analysis of DEGs

Gene Ontology (GO) is an online database providing bioinformatic information relating to gene and biological functions at different levels ranging from molecular interactions to organized tissues [12, 36]. Kyoto Encyclopedia of Genes and Genomes (KEGG) is a database containing high-level functionalities of the biological systems at different scales. We utilized WEB-based GEne SeT AnaLysis Toolkit, a platform supported by abundant bioinformatic data (http://webgestalt.org/), to conduct GO functional enrichment and KEGG pathway enrichment analysis. GO and KEGG analysis was applied for up-regulated genes (n= 7,147) and down-regulated genes (n= 6,528) respectively. GO analysis of biological pathways, cellular component, and molecular functions were applied to the DEGs that have selected from previous screening. Therefore, we identified the biological features shared by primary tumor cells of glioblastoma, and the two groups of concurrently regulated genes were responsible for were identified. A more comprehensive investigation into the cellular pathway from mRNA expression to the performance of GBM cells could be attained by identifying KEGG pathways involving the DEGs and thus reconstructing a system of specific metabolic pathways and gene interactions that contributed to the developmental progress of GBM.

### Construction of Protein-protein Interaction (PPI) network

The web-based search tool STRING (https://string-db.org/), which can retreat interacting genes, facilitated the manipulation of inputted protein data to visualize the relationship between our speculated vital proteins. STRING is utilized to map the DEGs and detect possible relationships among them. A down-stream PPI network analysis was therefore conducted for the proteins encoded by up- and down-regulated genes extracted from the result of previous analytical procedures. The interaction sources for the PPI network were selected as displaying Experiments and Databases with an interaction score of 0.700 as the threshold.

### Quantitative Real-Time Polymerase Chain Reaction (qRT-PCR)

The qRT-PCR was used to detect the expression of FPR2, VEGFA, SERPINA1, SOX2, PBK, ITGB3 in human glioma cell lines, compared with HEB cells. Total RNA was extracted from cells. Then the RNA was reversely transcribed to cDNA by using a Reverse Transcription Kit, and then the synthesized cDNA underwent qPCR according to the manufacturer’s protocol. The reactions were performed on the QuantStudio Dx instrument. The relative gene expression levels were calculated using the GAPDH.

### SDS-PAGE and Western blots

The cells were harvested and lysed in a lysing buffer. 10 – 20 μL of each sample per lane were added separated on sodium dodecyl sulfate-polyacrylamide gelelectrophoresis (SDS-PAGE). The proteins were immobilizedand transferred to a polyvinylidene difluoride (PVDF) membrane (1). After blocking the non-specific proteins on membrane with 5% fat-free milk and 0.1% Tween-20 in tris-buffered saline with Tween (TBST) for 1.5 h, primary antibody, diluted in 1:1000, were used against FPR2. PVDF membrane was incubated with the primary antibodies overnight at 4 °C. Anti-rabbit secondary antibody conjugated with horseradish peroxidase (1:1000) were added at room temperature for 1h.

## RESULTS

### Screening of DEGs in GBM and normal brain tissues

A total of 11 GBM and 5 normal gene expression profiles from TCGA were used in this study to identify the DEGs between GBM patient brain tissues and normal brain tissues. We detected a total of 13,675 differential expressed genes (7,147 up-regulated and 6,528 down-regulated genes) based on selection criteria. In addition, FPR2, VEGFA, SERPINA1, SOX2, PBK, and ITGB3 were identified because these six genes are presented as highly up-regulated (**Table 1**). GBM and normal samples were screened for DEGs through the construction of volcano plots (**Figure 1a**), Pearson’s correlation plot (**Figure 1b**), and heat map (**Figure 1c**) using the R package. A heat map of the up-and down-regulated DEGs were drawn using t, which could clearly discriminate GBM and normal brain tissues and show the general expression pattern of the selected DEGs.

**Table 1.**
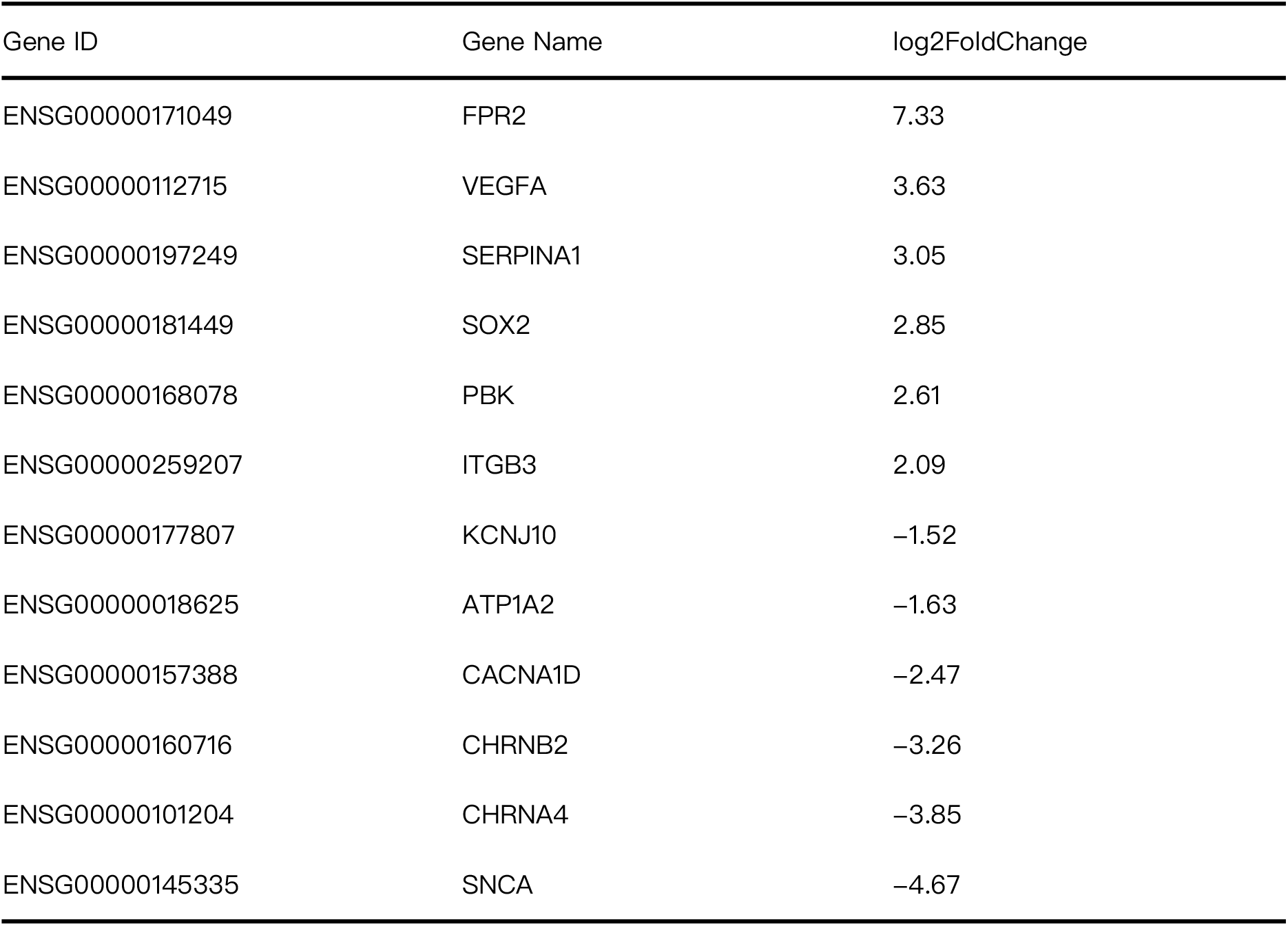
DEGs of Interest. Expression level of 12 selected genes significantly up- or down-regulated in GBM primary tumors based on analysis of TCGA-GBM samples.

**Figure 1.**
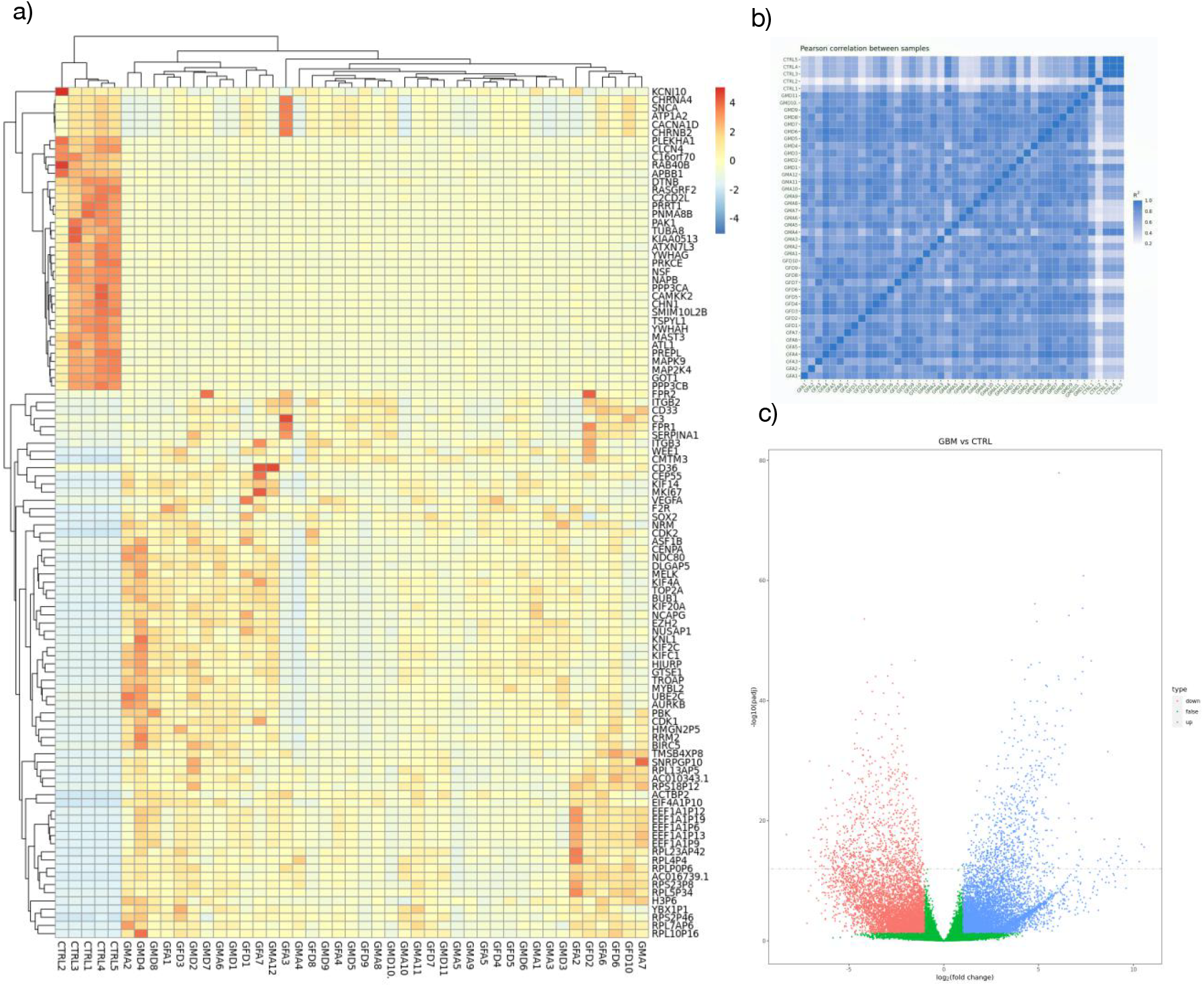
Differential gene expression heatmap and volcano plot between GBM and Control samples from TCGA gene expression profiles. **a)** Heatmap of up-regulated DEGs shows the general distribution of 104 significant DEGs of up-regulation and the down-regulation. **b)** Pearson’s correlation coefficient plot for DEGs screening. **c)** Volcano plots showing the log2FoldChange of DEGs. The red nodes represent the down-regulated DEGs, and the blue nodes represent the up-regulated. The green region refers to DEGs that are considered as insignificant.

### GO and KEGG analysis of up-regulated and down-regulated genes

To further explore the selected DEGs, WebGestalt was used to obtain the results of GO functions and KEGG pathway enrichment analysis. All the up-regulated and down-regulated DEGs were imported to the WebGestalt website (**Figure 2a**). GO analysis results demonstrated that up-regulated and down-regulated DEGs were particularly enriched in the following biological processes (BP): response to stress, metabolic processes, and response to a stimulus for up-regulated DEGs, which are all vital immune process and plays a crucial part in detecting and eliminating abnormal cells, protein and ion binding, nucleic acid binding, and hydrolyze activity for down-regulated DEGs. Moreover, GO cellular component (CC) analysis showed that both up-regulated and down-regulated DEGs were enriched in the membrane, nucleus, and membrane-enclosed lumen; whereas the down-regulated DEGs enriched in the plasma membrane and cell periphery.

**Figure 2.**
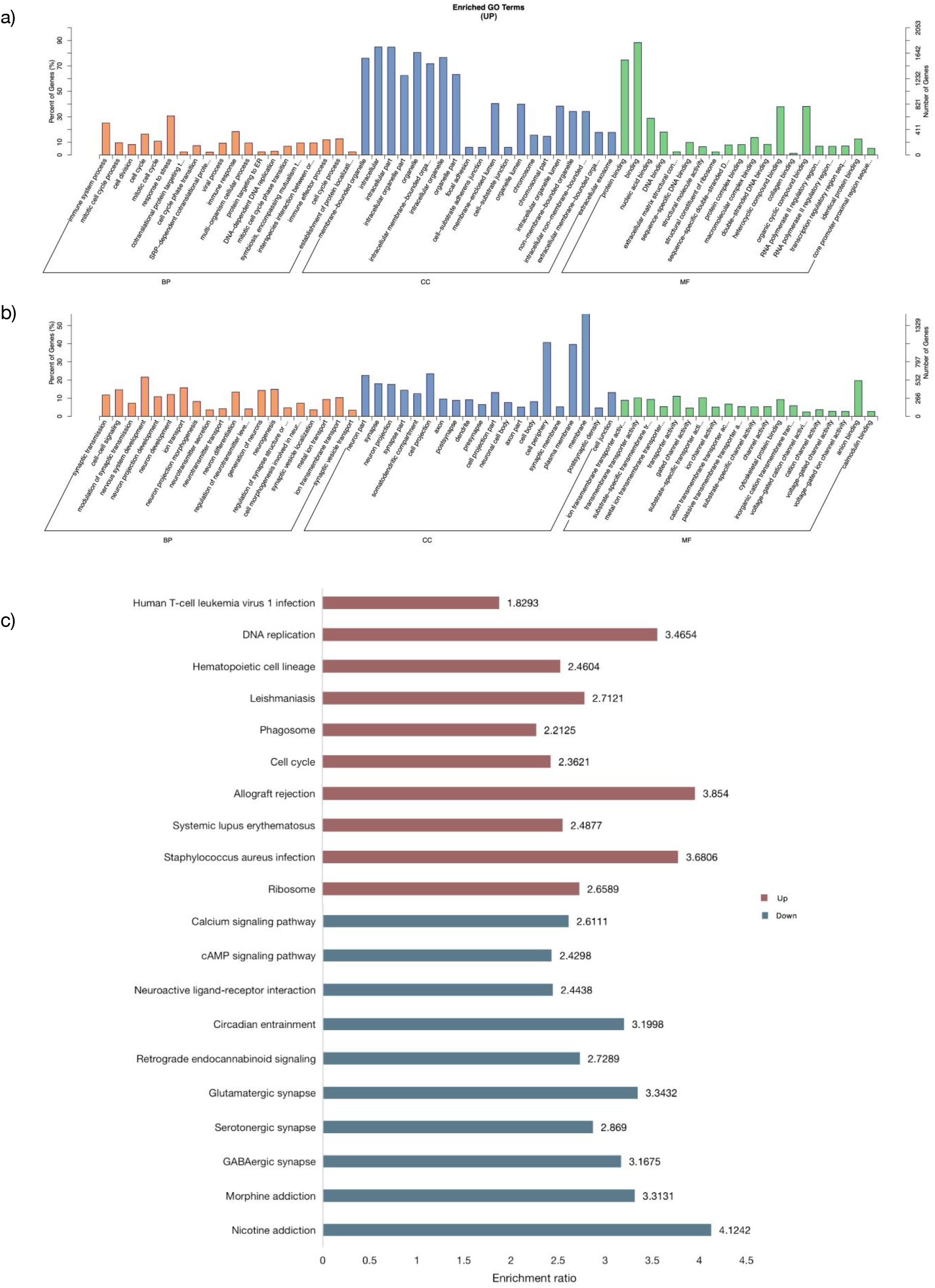
GO and KEGG analysis results of DEGs associated with GBM and normal brain tissues. **a)** Enriched GO terms of up-regulated genes. **b)** Enriched GO terms of down-regulated genes. **c)** Most significantly enriched KEGG pathway of the up-regulated DEGs. GO refers to gene ontology; BP for biological process; MF for molecular function; CC for cell component; KEGG for Kyoto Encyclopedia of Genes and Genomes.

The most significantly enriched KEGG pathways (**Figure 2b**) for up-regulated included Allograft rejection, Staphylococcus aureus infection, DNA replication, and down-regulated genes, which included Nicotine addiction, Glutamatergic synapse, and Circadian entrainment. The result derived from our analysis corresponded with the previous investigation [6, 14, 19, 38] in that the GO functions and KEGG pathways highlighted in this study composed mainly participated in inflammatory mechanisms, cell signaling, and cell division that played an indispensable role in cancer progression [29].

Protein-protein interaction network construction and identification of hub genes The most enriched genes in biological functions and pathways related to GBM development were selected and 94 up-regulated genes from the upstream functional analysis were used to formulate a PPI network that was displayed (**Figure 3**). In this investigation, particular interest was placed in the selected up-regulated DEGs including FPR2 and ITGB3. These genes played a significant role in the biological pathway leading to GBM progression, as a result, our ensuing research was focused on investigating the interrelationship between these key genes and tumor activity in GBM patients.

**Figure 3.**
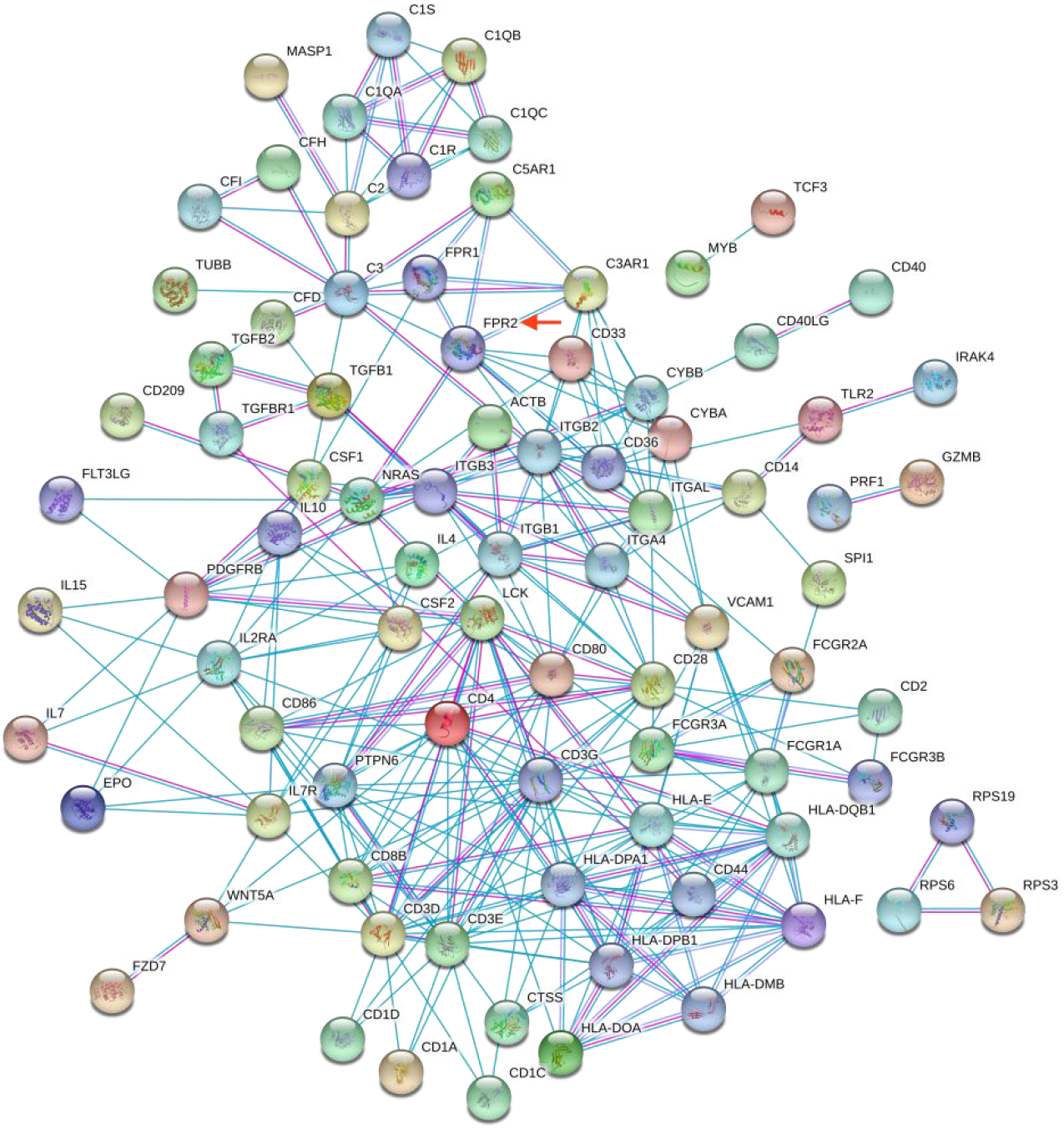
Protein interaction network constructed from chosen significant up-regulated DEGs. FPR2 indicated with a red arrow in the diagram which was identified as a hub gene in the up-regulated PPI network.

### Six hub genes expression in RT-qPCR

The RT-qPCR analysis was done to detect the expressions of 6 hub genes (FPR2, VEGFA, SERPINA1, SOX2, PBK, ITGB3) (**Supplementary 1**) to validate the results in the bioinformatics analysis. The expression of all of the 6 genes was significantly increased in glioma cell lines compared to a different extent compared to the HEB cell line (**Figure 4**), which further verify the differential analysis results [41]. Among these investigated hub genes, FPR2 has the highest expression levels. These results revealed that FPR2 may contribute to the progression of glioma.

**Figure 4.**
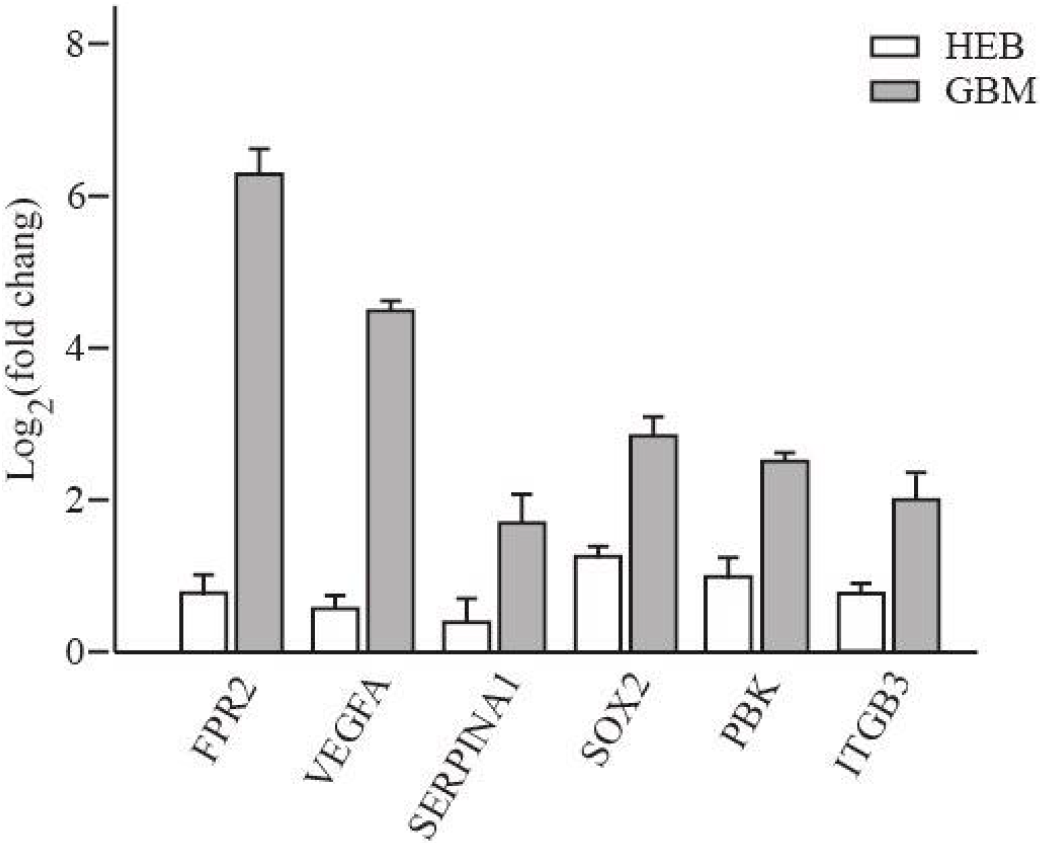
Real-time polymerase chain reaction shows the expression level of FPR2, VEGFA, SERPINA1, SOX2, PBK, ITGB3. The mRNA level of the 6 genes in the glioma samples were detected. All of them have shown considerable increase in the glioma cell lines, especially FPR2. HEB refers to normal brain glia cells.

### Western blotting analysis shows the up-regulation of FPR2 in glioma cells

To examine the expression level of FPR2 in GBM and NBT, we performed Western blotting to demonstrate the differences. Over-expression of FPR2 was observed in glioma cell lines, which shows high concordance to both differential analysis and qPCR results (**Figure 5**).

**Figure 5.**
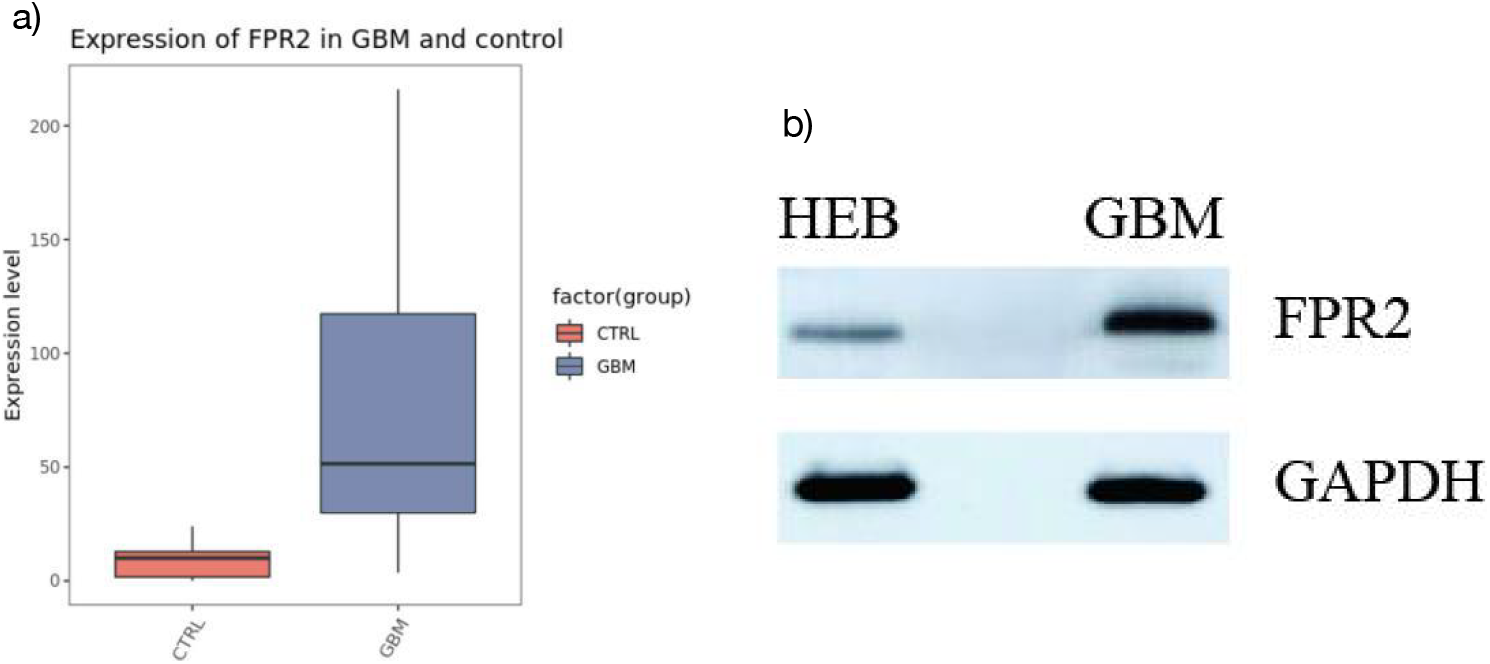
Comparison between the control and GBM group. **a)** Boxplot showing over-expression of FPR2 in GBM patients compared to Control group. **b)**Representative western blots showing higher FPR2 expression in glioma patients to normal brain cells. HEB refers to normal brain glia cells.

## DISCUSSION

Due to the invasive growth and high lethality [13, 33], the overall survival and prognosis of glioma are poor [1]. Therefore, effective biomarkers for evaluating the progression of glioma are important to the treatment and prognosis of patients.

In this study, we aimed to identify the differentially expressed genes in glioblastoma multiforme and normal control tissue, which could be used as potential targets for glioma therapy or prediction for the cancer progression [41, 45]. We identified 7,147 up-regulated genes and 6,528 differently expressed genes that were shared between GBM and Control Group. The results were screened by the criteria: |logFC| > 2, FDR<0.05. Among these DEGs, we selected 6 hub genes (FPR2, VEGFA, SERPINA1, SOX2, PBK, ITGB3) with a higher degree of connectivity and could be a potential biomarker for GBM prognosis [20, 27, 47, 49]. The considerably different expression level of FPR2 was presented in violin plots and box plots.

Formyl peptide receptor 2 (FPR2) is classified as the formyl peptide receptor (FPR) family subordinated to the GPCR superfamily. FPR2 as well as FPR1 participate in multiple intracellular signaling pathways ranging from activation of various protein kinases and phosphatase to tyrosine kinase receptors transactivation [16, 17, 21, 23]. The significance of these crucial cellular pathways within cells also suggested that any dysfunction of the activated pathways could lead to disorder within the body [47]. FPR2 was also known as FPR2A, FPRH1, FPRH2, and FPRL1 which was a key gene previously identified as a promotor of several cancers including colorectal cancer [25], gastric cancer, ovarian cancer, breast cancer [18], lung cancer, and glioblastoma. Former research indicated that FPR2 expression was also detected in normal glial cells and brain tissues but significantly higher in glioma cell lines and glioma tissues [22]. Besides, silencing of FPR2 in glioma cell lines had been proved to inhibit growth, invasion, and migration but promote the apoptosis of tumor cells, whose mechanisms might be associated with the inhibitory expression of cyclin D1 and VEGF.

Proven by the GO and KEGG analysis, the regulation of inflammatory reactions by the FPRs family is indispensable in the immune system and also interact with the Complement receptors 3 (CR3) [10]. “Compound 43” was discovered as an FPRs agonist and the agonists can induce a respiratory burst in leukocytes and activated C3 receptor (CR3) mobilization to the surface of the cell. Moreover, when high levels in CR3+ mononuclear phagocytes presents, FPR2 gene expression also increases [31].

The biological process and functions of these genes were evaluated by GO analysis and KEGG pathway enrichment analysis. In the KEGG analysis, DEGs were concentrated in Hematopoietic cell lineage, Complement, and coagulation cascades, and Cytokine-cytokine receptor interaction (CRI). Importantly, FPR2 and FPR1 has been repeatedly reported involved in ligand recognition and signaling in the CRI [42], with FPR1 being identified as enhancing motility and invading ability [48]. The over-expression of FPR2 could resolve inflammation and proliferation in astrocytomas [11, 34], which was consistent with our results. Thus, we believe FPR2 could be a novel biomarker of GBM and predict the prognosis of GBM patients. And to our knowledge, this is the first study that comprehensively examines the FPR2 gene with associations to GBM and identifying it as a viable target for GBM treatment. The mechanism of how the FPR2 gene regulates GBM requires further research.

Furthermore, the construction of the up-regulated PPI and down-regulated PPI networks between regulators and related genes was explored, which lucidly demonstrated the interactions between DEGs in GBM that might have a profound effect on the progression of the glioblastoma. FPR2 was validated as a critical gene that is related to glioma biological processes [43]. Highly relevant nodes, VEGFA which have been claimed to have a significant effect in tumor proliferation, invasiveness, migration, and angiogenesis [2], mainly operate in the activation of quiescent endothelial cells and promoting cell migration and proliferation [28]. Within the same family, studies also have shown that highly malignant glioma will selectively express a high level of FPR1, which also plays a role in promoting tumor progression. FPR1 has also been reported in many non-hematopoietic cells, such as colon epithelial cells, lung, and hepatocytes cells [23].

Although there are important discoveries revealed by this study, there are also limitations. First, the size of the control group used in this study was limited and was inadequate to appropriately represent a larger population. The number of cases utilized needs to be expanded to obtain more accurate results. Second, this investigation conducted laboratory-based experiments using purchased GBM cell lines instead of patient-derived pathological samples. It would be necessary to use biopsy samples and include animal experiments for a more comprehensive understanding of the molecular mechanism behind GBM progression. Moreover, the precise molecular mechanism and detailed regulatory networks involved in the occurrence and development of GBM need to be further explored by conducting subsequent investigations.

Current medical and therapeutical options of GBM have attained limited success, yet genomic profiling analysis has presented comprehensive solutions to analyze gene data in large quantities to associate prognosis outcome, tumor metastasis and formation, and other clinical features to patient survival [46]. Overall, we concluded that the FPR2 gene can be used as a suitable prognostic biomarker for GBM and the findings in this research have presented its significant potential, that with more attentive and specific explorations in the future, it will expedite further development of possible treatments.

## Supplementary

**1.**
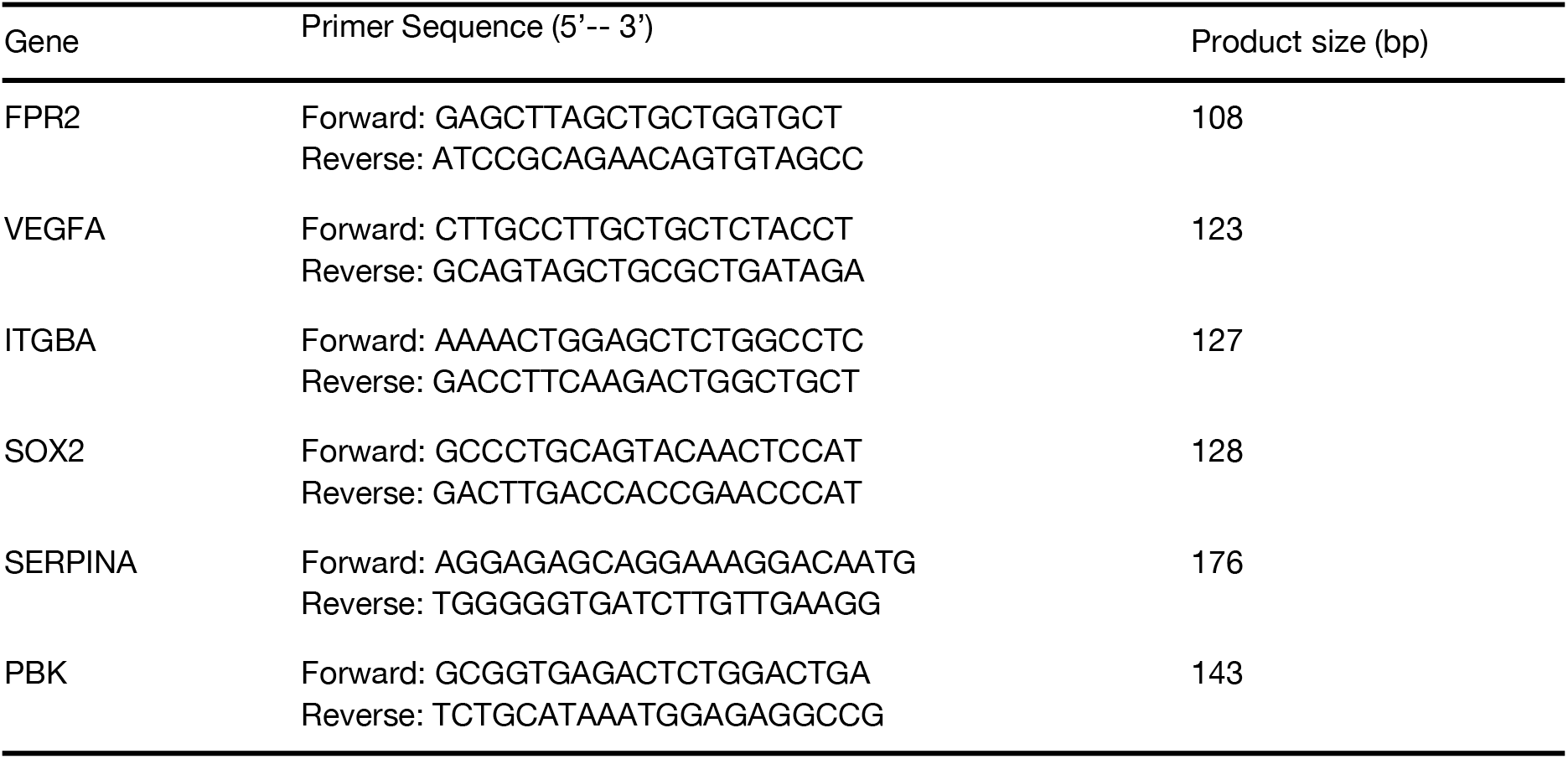
Real time qPCR primer sequences used in this experiment. qPCR was performed for six selected up-regulated genes: FPR2, VEGFA, ITGBA, SOX2, SERPINA, and PBK.

## Notes

### Competing Interest Statement

The authors have declared no competing interest.

